# Phenotypic plasticity explains apparent reverse evolution of fat synthesis in parasitoid insects

**DOI:** 10.1101/2020.07.17.208876

**Authors:** Bertanne Visser, Hans T. Alborn, Suzon Rondeaux, Manon Haillot, Thierry Hance, Darren Rebar, Jana M. Riederer, Stefano Tiso, Timo J.B. van Eldijk, Franz J. Weissing, Caroline M. Nieberding

**Affiliations:** Evolution and Ecophysiology group, Biodiversity Research Centre, Earth and Life Institute, UCLouvain, Croix du Sud 4-5, 1348 Louvain-la-Neuve, Belgium.; Chemistry Research Unit, Center of Medical, Agricultural, and Veterinary Entomology, Agricultural Research Service, United States Department of Agriculture, 1600 SW 23rd Drive, Gainesville, FL 32608, USA.; Ecology of Interactions and Biological Control group, Biodiversity Research Centre, Earth and Life Institute, UCLouvain, Croix du Sud 4-5, 1348 Louvain-la-Neuve, Belgium.; Department of Biological Sciences, Emporia State University, 1 Kellogg Circle, Campus Box 4050, Emporia, KS 66801, USA.; Groningen Institute of Evolutionary Life Sciences, University of Groningen, Nijenborgh 7, 9747 AG Groningen, the Netherlands.; Evolutionary Ecology and Genetics group, Biodiversity Research Centre, Earth and Life Institute, UCLouvain, Croix du Sud 4-5, 1348 Louvain-la-Neuve, Belgium.

## Abstract

Over the last few decades, numerous examples have been described where a trait that was once lost during the course of evolution had been regained. Here, we argue that such reverse evolution can also become apparent when trait expression is plastic in response to the environment. We tested this hypothesis for the loss and regain of fat synthesis in parasitic wasps. Wasps from lineages that supposedly regained lipogenic ability ~80 million years ago were grown under a fat-poor or fat-rich environment. In line with our hypothesis, it turned out that fat synthesis had not been lost and regained, but was only switched on in low-fat environments. Functional protein domains of key lipogenesis genes were also found in other parasitoid species, suggesting that plasticity of fat synthesis may be more widespread in parasitoids. Individual-based simulations then revealed that a switch for plastic expression can remain functional in the genome for thousands of generations, even if it is only used sporadically. The evolution of plasticity may thus also explain other examples of apparent reverse evolution.

## Introduction

There are numerous cases where a complex adaptation has been lost in the evolutionary history of a lineage^1^, e.g., legs in snakes, teeth in birds, and the ability to fly in ratites. If a trait is of no use for an extended period of time, it can be selected against and/or decay by genetic drift and the accumulation of deleterious mutations^2^. The last decades have seen a surge of papers reporting reverse evolution, i.e., cases in which a trait that was once lost had reappeared^3,4^. Highly cited studies include floral adaptations^5,6^, reproductive/breeding systems^7–10^, and anatomical changes^11–13^. Trait regain over extensive periods of time is an evolutionary puzzle, because mutation accumulation in underlying genetic pathways makes the re-evolution of functional activity by reverse mutations highly unlikely^14^.

Here, we scrutinize a reported case of reverse evolution: the apparent loss and regain of an essential metabolic trait, fat synthesis^15,16^. Fat is synthesized when a surplus of sugars (and other carbohydrates) is available in the diet^17^, providing an energy reserve for future use. Fat is critical for survival and reproduction in nearly all living organisms; hence underlying metabolic and genetic pathways for fat synthesis are typically highly conserved from bacteria to humans^18–21^. A comparative study in 2010 revealed that parasitic wasps (i.e., hymenopteran parasitoids) lost the ability to synthesize fat in their common ancestor more than 200 million years ago^16,22^. The loss of fat synthesis was thought to result from consumption of host lipids, because parasitic wasps feed on another insect to complete their own development^23^. On at least three separate occasions, fat synthesis re-appeared throughout the wasp phylogeny, particularly in species with wide host ranges (i.e., generalists)^16^. Host availability for generalists is expected to be stochastic; hence constitutive fat synthesis would be critical for their survival and reproduction.

Recently, active fat synthesis was also found in several closely related wasps in the genus *Nasonia*^24^, thought to have lost the ability for fat synthesis^16,25^. The observed variation in lipogenic ability could have resulted from genetic evolution due to local adaptation. A generalist species, thought to have regained fat synthesis in the 2010 study, *Leptopilina heterotoma*, was the subject of more detailed work that hinted at an alternative hypothesis to that of genetic evolution: Some field-collected populations synthesized fat, while others did not^26^. When these experiments were replicated on a larger scale, none of the populations synthesized fat^26^. These contradictory results between studies could be explained by plasticity-based local adaptation, where the environmental cue used for the expression of fat synthesis is the fat content of the host. Indeed, in the first experiment wasps emerged with almost half the percentage of fat (i.e., ~16%) compared to wasps that emerged from the second experiment (i.e., ~28%)^26^. The host strain used for the first experiment was thus thought to have been much leaner compared to the strain used for the second experiment, inducing lipogenesis at least in some populations.

Considering the time required for mutational changes to appear and to be maintained, we hypothesized that fat synthesis in parasitic wasps, and other parasitoids, was not lost and regained due to mutational changes in the metabolic pathway (genetic evolution hypothesis), but rather that fat synthesis shows plastic expression (on or completely off) in response to the local environment (adaptive plasticity hypothesis). Here, we present support for the plasticity-based hypothesis with three distinct lines of evidence: *1)* experimental evidence that the wasp *L. heterotoma* shows plasticity of fat synthesis (sisters can switch fat synthesis on or completely off); *2)* a comparative DNA sequence analysis that suggests key enzymes involved in fat synthesis are still functional in various parasitoid insects, including a parasitic fly and beetle, and *3)* individual-based simulations that show that a switch for plasticity remains functional in the genome over thousands of generations, even when only used sporadically.

## Results

### A family-based experimental design reveals that fat synthesis is plastic

We first tested whether wasps would switch lipogenesis on when development occurred on a lean host. To this end, we first let females from four field-caught populations of *L. heterotoma* develop on two naturally co-occurring host species: fat-poor (“lean”) *Drosophila simulans* and fat-rich (“fat”) *D. melanogaster* (containing 63 ± 3 μg and 91 ± 4 μg, mean ± 1SE storage fat, respectively; F_1,17_ = 35.95; p < 0.0001). When wasps developed on lean *D. simulans*, fat synthesis had occurred in 3 out of 4 wasp populations (Table 1). In contrast, wasps did not significantly increase their fat content when developing on fat *D. melanogaster*. These findings confirm previous findings^26^, and suggest that wasp fat synthesis depends on the host environment.

**Table 1:**
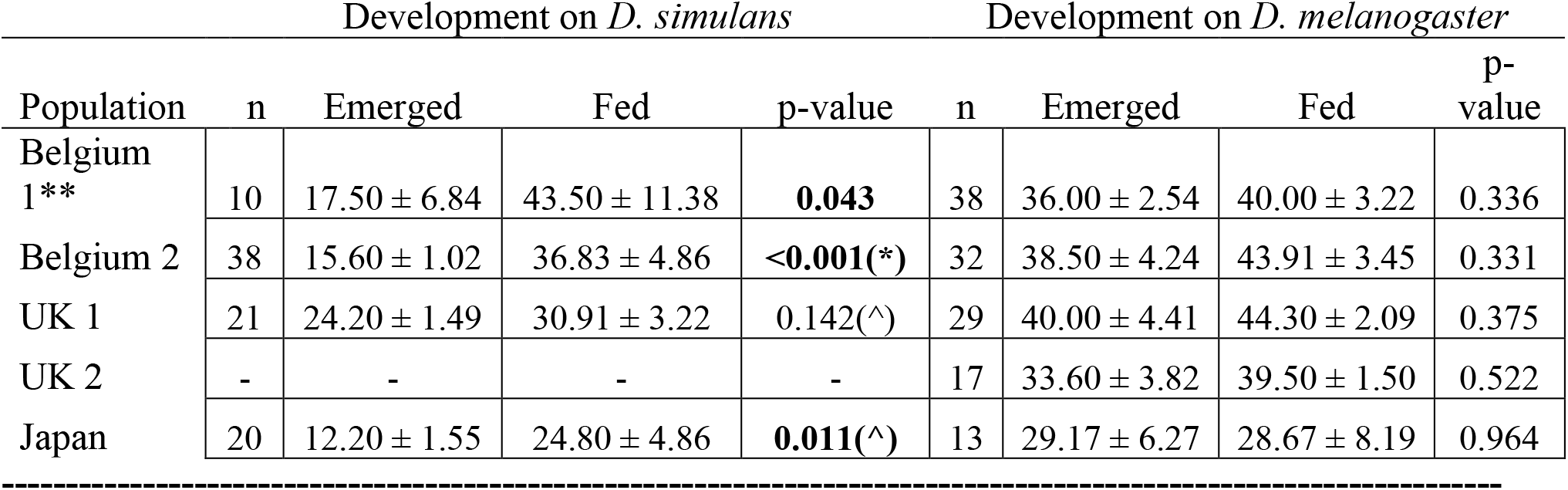
Wasps synthesize fat in a fat-poor environment. Mean absolute fat amount ± 1se (in μg) was quantified in adult wasps from field-caught *L. heterotoma* populations raised on two hosts (lean *D. simulans*, left part of the table; fat *D. melanogaster*, right part of the table) and at two time points during adult life (Emerged: just after emergence; Fed: having fed for 7 days after emergence). P-values reveal whether 7 days of feeding led to a significant increase in fat content, indicating that fat synthesis had occurred. Three of the four populations tested on *D. simulans* exhibited fat synthesis on the lean host, but no fat synthesis on the fat host. T-tests were performed when data was normally distributed and variances equal with (^) or without log transformation. The non-parametric Mann-Whitney U test was used for non-normal data or data with unequal variances (*). Population UK2 was not available for testing on *D. simulans.*

The population-level comparison of wasp fat content described above is a common, but crude measure that not always detects the occurrence of fat synthesis reliably. Even in case of active fat synthesis, fat content can stay constant or even decrease if, for example, fats are burned at a faster rate than at which they are produced^27^. This means that equal or decreasing fat amounts do not conclusively indicate a lack of lipogenesis. To unequivocally demonstrate that fat synthesis can be induced plastically, we turned to stable isotope tracing followed by GC-MS (Gas Chromatography-Mass Spectrometry) analyses^28,29^. Incorporation of stable isotopes after feeding depends on fat synthesis and is traced directly into the fatty acid fraction. A significant increase in stable isotope levels compared to controls (without access to stable isotopes) thus demonstrates active fat synthesis, even when lipids are burned. We used a split-brood family design where daughters of a single mother (sharing 75% of their genome) were allowed to develop on either lean *D. simulans* or fat *D. melanogaster*. Seventeen families, belonging to five field-caught populations, showed a (much) higher fat metabolism in the fat-poor environment (*D. simulans*) than in the fat-rich environment (*D. melanogaster*) (Fig. 1). These results confirm that fat synthesis is indeed a plastic trait that is induced in response to low host fat content and completely shut off in response to high host fat content. Notice also that the 17 families strongly differ in their environmental response, both in their baseline level of fat synthesis (on fat *D. melanogaster*) and in the slopes of their reaction norms. We thus provide conclusive evidence that fat synthesis is plastic in *L. heterotoma*.

**Fig. 1:**
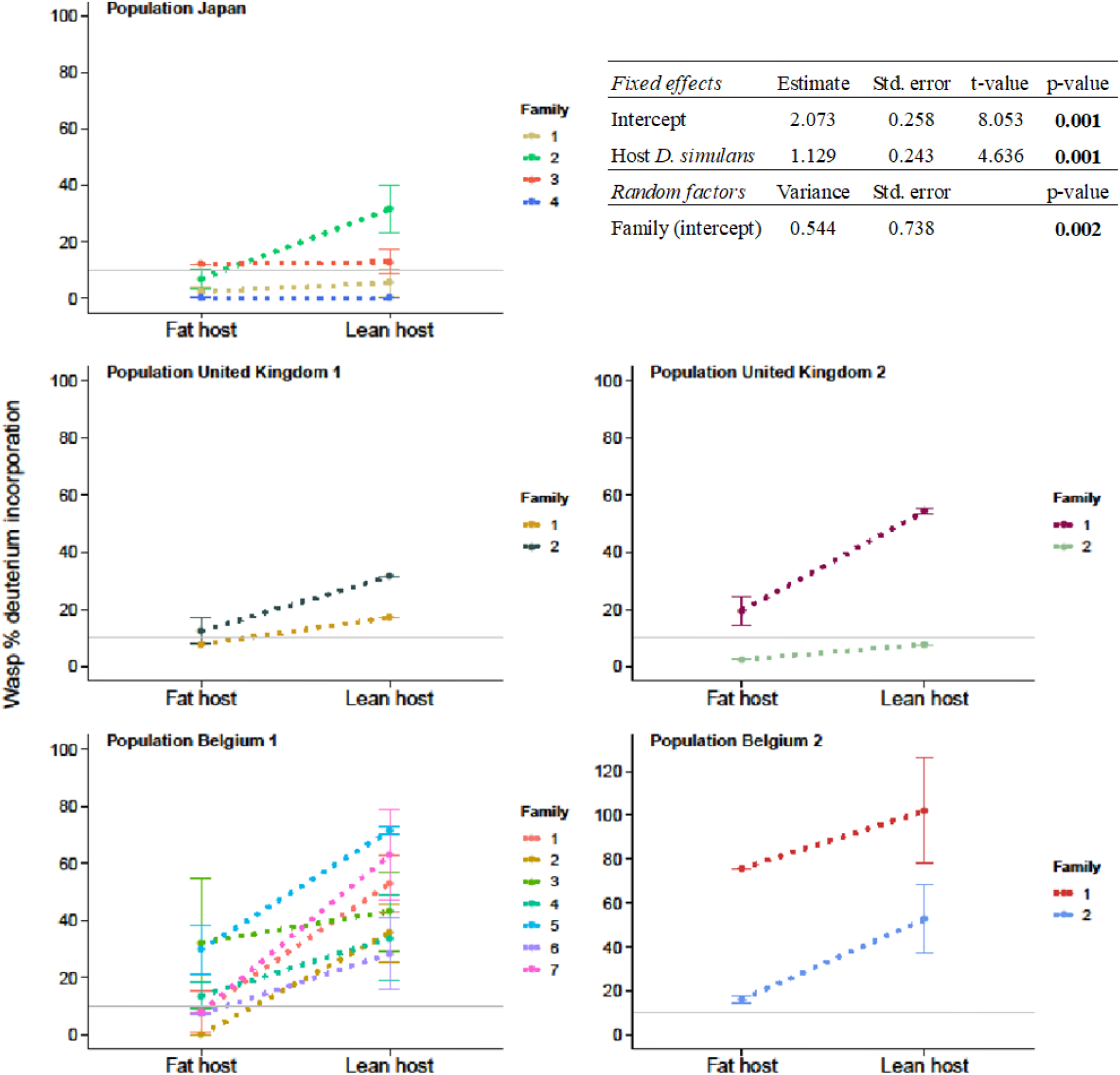
Phenotypic plasticity in five field-caught wasp populations. Incorporation of stable isotopes into the fatty acid fraction of offspring from 17 families developing in a fat-rich environment (fat host *Drosophila melanogaster,* left in each graph) and in a fat-poor environment (lean host *D. simulans,* right in each graph; n = 138). The horizontal gray line indicates that a stable isotope incorporation below 10% is considered insufficient evidence for the occurrence of fat synthesis.

To rule out the possibility that the above results reflect a host species effect, rather than an effect of the host’s fat content, we repeated the experiment reported in Table 1, but now testing fat synthesis using a single host species, *D. melanogaster*. By reducing the sugar content in the diet of *D. melanogaster*, we were able to generate leaner flies (i.e., pupae containing 52 ± 3 μg storage lipids, mean ± 1SE, compared to 91 ± 4 μg storage lipids, mean ± 1SE; F_1,22_ = 71.18, p < 0.0001). In line with findings for *D. simulans,* three out of four wasp populations synthesized fat on lean *D. melanogaster* hosts (Table 2). We thus conclude that plastic fat synthesis is induced by host fat content, rather than other traits differing between *D. melanogaster* and *D. simulans.*

**Table 2:** Plasticity irrespective of host species identity. Mean absolute fat amount ± 1se (in μg) was quantified in adult wasps from field-caught *L. heterotoma* populations raised on fat-poor *D. melanogaster* at two time points during adult life (Emerged: just after emergence; Fed: having fed for 7 days after emergence). P-values reveal that fat synthesis took place in three of the four wasp populations, meaning that lipogenesis is also plastic when measured using the same host strain. T-tests were performed when data was normally distributed and variances equal with (^) or without log transformation. Population Belgium 1 was not available for testing on lean *D. melanogaster*.

| Population | Sample size | Emerged | Fed | p-value |
| --- | --- | --- | --- | --- |
| Belgium 2 | 31 | $27.40 \pm 1.82$ | $38.08 \pm 3.50$ | **0.018 (^)** |
| UK 1 | 33 | $25.09 \pm 2.51$ | $40.50 \pm 2.66$ | **<0.001** |
| UK 2 | 35 | $27.25 \pm 2.60$ | $38.62 \pm 2.70$ | **0.006** |
| Japan | 31 | $34.70 \pm 2.72$ | $34.36 \pm 3.69$ | 0.954 |

### Protein domains of key lipogenic genes are functional in parasitoid wasps, a beetle and a fly

The ability to synthesize fat when being placed in a low-fat environment indicates that key genes for fat synthesis have not lost their functionality in the *Leptopilina* genus^26,30^. Making use of the fact that the genetic molecular pathway underlying fatty acid synthesis is highly conserved across animal taxa^18–21^, we conducted a comparative analysis of coding sequences of acetyl coenzyme A carboxylase (ACC)^31^ and fatty acid synthase (FAS)^20^, two enzymes that are critical for the production of fatty acids, the raw materials for stored fat. We used the *acc* and *fas* gene coding sequences of *D. melanogaster* as a starting point, because this fly readily synthesizes fat^32,33^. Similar gene sequences were indeed found in the genome of *L. clavipes,* a sister species of *L. heterotoma,* and all functional domains of ACC and FAS enzymes were recovered, suggesting fully functional coding sequences in the *L. clavipes* genome (Fig. 2). We then expanded our search for *acc* and *fas* functional coding sequences and protein domains to more distantly related parasitoids presumed to have lost fat synthesis independently^16^: the hymenopteran *Goniozus legneri* (family Bethylidae), the dipteran *Paykullia maculata* (family Rhinophoridae), and the coleopteran *Aleochara bilineata* (family Staphilinidae)(Fig. 2). No stop codons were found in any of the protein domains and ACC and FAS amino acid sequences of all these species aligned (Supplementary Texts 1 and 2), suggesting that these two critical genes for fat synthesis have been conserved throughout the repeated evolution of parasitism in insects. Further tests on these and other parasitoids are now needed to confirm plasticity of fat synthesis at the phenotypic level, but emerging results in other parasitoid systems, e.g., *Nasonia* species^24^ and *Meteorus pulchricornis*^34^ strengthen the notion that plasticity of fat synthesis may be more widespread in parasitoids.

**Fig. 2:**
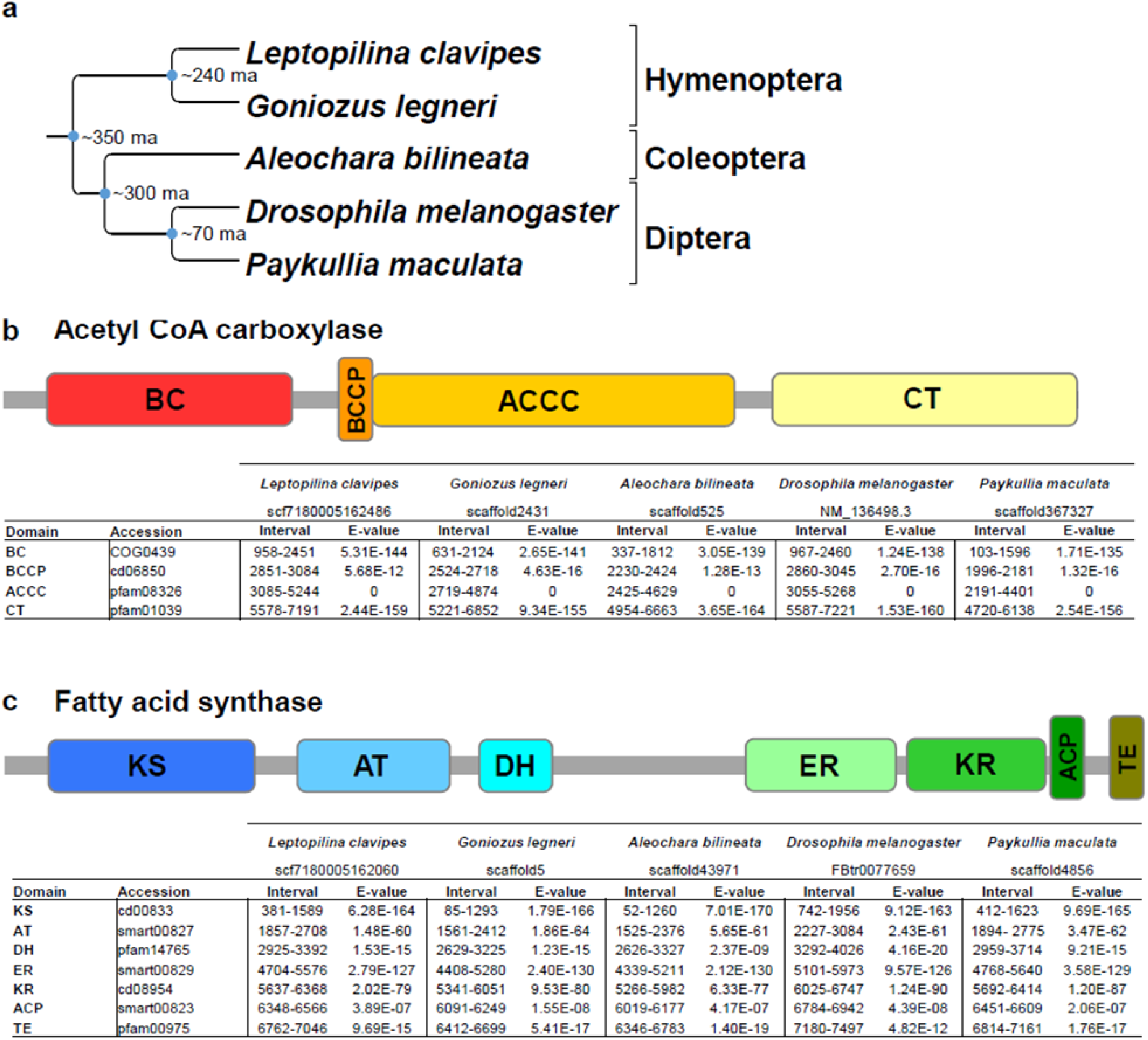
Conservation of two genes crucial for fatty acid synthesis in four parasi toid insects that supposedly had lost lipogenic activity. Long evolutionary divergence times (up to 350 MA) separate the insect *Drosophila melanogaster* (that synthesizes lipids constitutively) and 4 parasitoid insects that were assumed to have lost the ability to synthesize lipids(l0) (A). Acetyl coenzyme A carbo xylase (ACC) and fatty acid synthase (FAS) are two essential genes for the production of fatty acid: the presence of all domains of ACC (B) and FAS (C) genes from *D. melanogaster* in the four parasitoid genomes reveals that the functional activity of the two genes is conserved in these insects. A table containing the detailed length and position of the different functional domains forming the two genes, as well as conservation level of the nucleotide sequence of the domain s (e-values; the lower the e-value, the higher the significance of the match) are shown for each species. Abbreviations: BC = Biotin carboxylase; BCCP = Biotin carboxyl canier protein; ACCC = Acetyl-coAcarboxylase central region; CT = Carboxyl transferase domain; KS = Ketoacyl synthase; AT = Acyl transferase; DH = Dehydratase; ER = Enoyl reductase; KR = Ketoacyl reductase; ACP = Acyl canier protein; TE = Thioesterase. Accession numbers refer to the conserved domain identifier on NCBI’s Conserved Domain Database. Parasitoid transcript identifiers are provided underneath each species name.

### A switch for plastic expression of adaptive traits is maintained in the genome when rarely used

The question then arises whether a switching device that is not used for extensive periods of time (more than 100 million years) should not be lost during the course of evolution. To investigate this, we ran individual-based simulations that monitored the sustained functionality of a switching device (a gene regulatory network that could decay by mutation) that is only sporadically used in evolutionary time. Fig. 3 shows that the switching device rapidly disintegrates (red simulations) if it is never used. However, even very infrequent use (pink: every 100 generations; purple: every 1000 generations) suffices to keep the switching device largely intact. Interestingly, the switching device does not erode gradually, but instead slowly evolves an improved performance over evolutionary time (i.e., the percentage of correct decisions increases with the increasing number of generations). An inspection of the evolving gene regulatory networks reveals that they become more and more robust (i.e., less and less affected by mutational decay), in line with earlier findings on network evolution^35^.

**Fig. 3:**
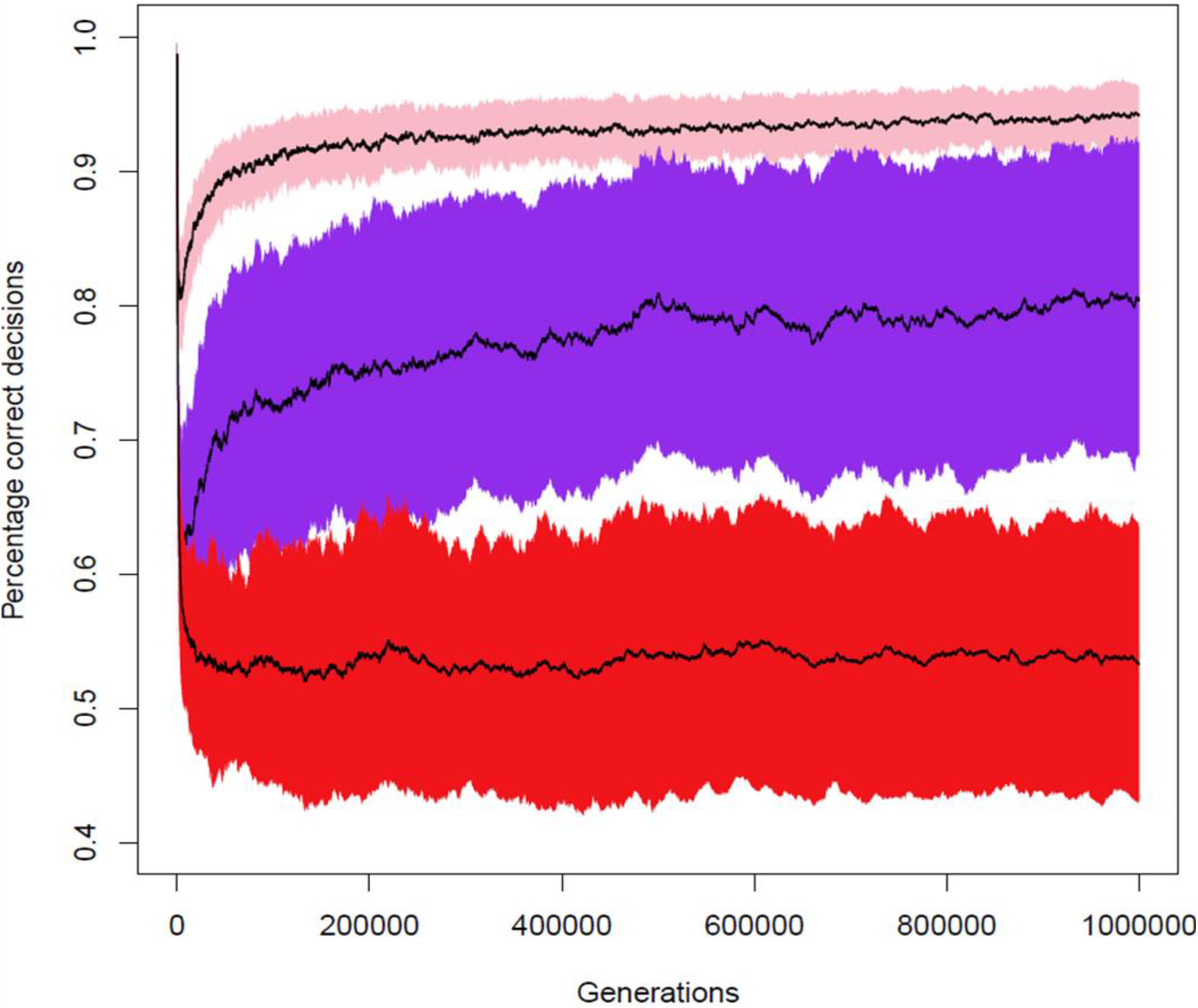
Sporadic activation is sufficient for the maintenance of adaptive plasticity. Long-term individual-based simulations showing how the performance of a gene-regulatory network (GRN) underlying adaptive plasticity changes in time when plasticity is only sporadically activated. We first evolved replicate GRNs in a variable environment where it is adaptive to switch on a metabolic pathway (fat synthesis) under low-fat conditions and to switch it off under high-fat conditions. In generation 0, a monomorphic population was established, where all N=l0,000 individuals were endowed with the same well-performing GRN (different across replicates). Subsequently, the population evolved subject to selection, mutation (μ=0.001 per gene locus) and genetic drift in a fat-rich enviromnent, where it is adaptive to constitutively switch off the metabolic pathway. Eve1y 100 generations, we monitored the performance of a representative sample of GRNs (percentage correct decisions) in the original (fat-variable) enviromnent: 1.0 means that the GRN is still making 100% adaptive decisions; 0.5 means that the GRN only makes 50% adaptive decision, as would be expected by a random GRN or a GRN that switches the pathway constitutively on or off. The coloured graphs show the average performance (± standard deviation) of the GRNs for three scenarios (100 replicates per simulation). Red: the population never again encounters the fat-variable enviromnent, leading to the loss of adaptive plasticity (convergence to 0.5). Pink: the individuals encounter a fat-variable enviromnent on average eve1y 100 generations; after an initial rapid drop in performance, a sustained high performance (>90% correct decisions) of the GRNs is regained after about 100,000 generations. Purple: the individuals encounter a fat-variable enviromnent on average every 1000 generations; after an initial rapid drop in performance, an intermediate performance (>75% correct decisions) is regained gradually.

## Discussion

We show that *L. heterotoma* does not represent a case of reverse evolution at the species/ population level (genetic evolution), but that fat synthesis is plastic (switched on or completely off) in response to the environment (adaptive plasticity). Large differences in slopes between families observed in Fig. 1 suggest that there is genetic variation for plastic expression of fat synthesis; hence adaptive plasticity in fat synthesis may itself evolve according to local fat content of host populations in the wild. The ability of these wasps to completely switch off fat synthesis, despite continued feeding on sugars, is unique and exceptional and we are unaware of a similar finding in other animals. A crucial pathway like fat metabolism is thus not constitutively expressed in parasitoids, as in other animals, but activated or deactivated in response to environmental conditions. This makes perfect sense, since they typically develop on fat-rich hosts that provide all the storage fat needed by the wasps. Yet, plasticity is required because there is considerable spatio-temporal variation in host availability and quality. *L. heterotoma* is a generalist wasp that can parasitize more than ten different *Drosophila* species that differ substantially in size and fat availability^36^. Moreover, there is considerable geographic and seasonal variation in host species diversity and community composition^37^. Hosts are further patchily distributed with overlapping generations, suggesting considerable spatial variation at a local scale^36^. *Drosophila* are further well known to show large variation in starvation resistance, which is typically correlated with fat content^38^. It is therefore likely that plasticity of wasp fat synthesis is adaptive and evolved in response to highly variable environmental conditions in host fat content. Future studies with natural populations should aim to provide the empirical evidence for genotype by environment interactions.

Previous documented cases of trait regain over long evolutionary time, in addition to the regaining of fat synthesis in parasitoids^16^, include the regaining of wings in stick insects^39^, the evolution of sexual reproduction from asexuality in mites^40^, among other examples^11^. These cases were all based on phylogenetic analyses. Such analyses were already shown to be problematic, because phylogenies do not necessarily provide a reliable representation of trait evolution^41–44^. Our results provide the first experimental evidence that macro-evolutionary patterns of trait reversals may in fact reflect trait plasticity: the trait is not “lost” or “regained” but is rather switched off or on, depending on environmental conditions. Intriguingly, such a regulatory switch can remain largely intact, even if it is only sporadically activated (Fig. 3). We consider it plausible that our findings are not restricted to fat metabolism in parasitoid wasps: the plastic regulation of trait expression could explain more cases of apparent trait loss and reappearance at macro-evolutionary time scales. Wing formation, for example, is often observed as an atavism (the sporadic occurrence of an ancestral phenotype) in otherwise wingless insects^45^, and wing polymorphism, i.e. plasticity in wing development, is common in insects in general^46^. Similarly, many asexual populations sporadically produce sexually reproducing individuals and plasticity in reproductive mode has evolved in several insect systems^47–49^. Hence, plasticity may be a common principle explaining apparent reverse evolution.

## Methods

### Experimental study and protein domain analysis

#### Insects

Hosts and parasitoids were maintained as previously described^26^. Five *Leptopilina heterotoma* (Hymenoptera: Figitidae) populations were used for experiments: a population from Japan (Sapporo), two populations from the United Kingdom (1: Whittlesford; 2: Great Shelford) and two populations from Belgium (1: Wilsele; 2: Eupen). Information on collection sites, including GPS coordinates, can be found in^26^.

#### Determination of host fat content

*D. simulans* and *D. melanogaster* hosts were allowed to lay eggs over 24 hours in glass flasks containing ~50mL standard medium^26^. After two days, developing larvae were sieved and ~200 were larvae placed in a *Drosophila* tube (ø x h 25×95; Dominique Dutscher) containing ~10mL medium. Seven days after egg laying, newly formed pupae were frozen at −18°C, after which fat content was determined as described in^26^, where dry weight before and after neutral fat extraction was used to calculate absolute fat amount (in μg) for each host. The host pupal stage was chosen for estimating fat content, because at this point the host ceases to feed, while the parasitoid starts consuming the entire host^36^. All data were analysed using R Project version 3.4.3^50^. Fat content of hosts was compared using a one-way ANOVA with host species as fixed factor.

#### Manipulating host fat content

To generate leaner *D. melanogaster* hosts, we adapted our standard food medium^26^ to contain 100 times less (0.5g) sugar per litre water. Manipulating sugar content did not alter the structure of the food medium, thus maintaining similar rearing conditions, with the exception of sugar content. Fat content of leaner and fatter *D. melanogaster* hosts was determined and analysed as described above.

#### Fat synthesis quantification of wasp populations

Mated female *L. heterotoma* were allowed to lay eggs on host fly larvae collected as described above with *ad libitum* access to honey as a food source until death. Honey consists of sugars and other carbohydrates that readily induce fat synthesis. After three weeks, adult offspring emergence was monitored daily and females were haphazardly placed in experimental treatments: emergence or feeding for 7 days on honey. Wasps were frozen at −18°C after completion of experiments. Fat content was determined as described above for hosts. The ability for fat synthesis was then determined by comparing fat levels of recently emerged and fed individuals, similar to procedures described in^16,26,28^. An increase in fat levels after feeding is indicative of active fat synthesis; equal or lower fat levels suggest fat synthesis did not take place. Each population was analysed separately (as in ^26^, because we are interested in the response of each population) for each host species using T-tests when data was normally distributed and variances equal, while the non-parametric Mann-Whitney U test was used for non-normal data or data with unequal variances.

#### Fat synthesis quantification using a familial design and GC-MS analyses

To tease apart the effect of wasp genotype and host environment, we used a split-brood design where the offspring of each mother developed on lean *D. simulans* or fat *D. melanogaster* hosts in two replicated experiments (experiment 1 and 2). In both experiments, mothers were allowed to lay eggs in ~200 2^nd^ to 3^rd^ instar host larvae of one species for four days, after which ~200 host larvae of the other species were offered during four days. The order in which host larvae were presented was randomized across families. Following offspring emergence, daughters were allocated into two treatment groups: a control where females were fed a mixture of honey and water (1:2 w/w) or a treatment group fed a mixture of honey and deuterated water (Sigma Aldrich)(1:2 w/w; stable isotope treatment) for 7 days. Samples were prepared for GC-MS as described in ^29^. Incorporation of up to three deuterium atoms can be detected, but percent incorporation is highest when only 1 deuterium atom is incorporated. As incorporation of a single atom unequivocally demonstrates active fat synthesis, we only analysed percent incorporation (in relation to the parent ion) for the abundance of the m+1 ion. Percent incorporation was determined for five fatty acids, C16:1 (palmitoleic acid), C16:0 (palmitate), C18:2 (linoleic acid), C18:1 (oleic acid), and C18:0 (stearic acid), and the internal standard C17:0 (margaric acid). Average percent incorporation for C17:0 was 19.4 (i.e. baseline incorporation of naturally occurring deuterium) and all values of the internal standard remained within 3 standard deviations of the mean (i.e. 1.6). Percent incorporation of control samples was subtracted from treatment sample values to correct for background levels of deuterium (i.e. only when more deuterium is incorporated in treatment compared to controls fatty acids are actively being synthesized). For statistical analyses, percent incorporation was first summed for C16:1, C16:0, C18:2, C18:1 and C18:0 to obtain overall incorporation levels, as saturated C16 and C18 fatty acids are direct products of the fatty acid synthesis pathway (that can subsequently be desaturated).

Data (presented in Fig. 1) was analysed by means of a linear mixed effects model (GLMM, lme4 package) with host (lean *D. simulans* and fat *D. melanogaster*) as fixed effect, population (Japan, United Kingdom 1 and 2, Belgium 1 and 2), family, and experiment (this experiment was conducted twice) as random factors, and percentage of incorporation of stable isotopes as dependent variable (n = 138). Non-significant terms (i.e. population and experiment) were sequentially removed from the model to obtain the minimal adequate model as reported in the table. When referring to “families,” we are referring to the comparison of daughters of singly inseminated females, which (in these haplodiploid insects) share 75% of their genome.

### Identification of functional *acc* and *fas* genes in distinct parasitoid species

To obtain *acc* and *fas* nucleotide sequences for *L. clavipes, G. legneri, P. maculata* and *A. bilineata*, we used *D. melanogaster* mRNA ACC transcript variant A (NM_136498.3 in Genbank) and FASN1-RA (FBtr0077659 in FlyBase) and blasted both sequences against transcripts of each parasitoid (using the blast function available at parasitoids.labs.vu.nl^51,52^. Each nucleotide sequence was then entered in the NCBI Conserved Domain database^53^ to determine the presence of all functional protein domains. All sequences were then translated using the Expasy translate tool (https://web.expasy.org/translate/), where the largest open reading frame was selected for further use and confirming no stop codons were present. Protein sequences were then aligned using MAFFT v. 7 to compare functional amino acid sequences between all species^54^.

### Modelling study

We consider the general situation where phenotypic plasticity is only sporadically adaptive and ask the question whether and under what circumstances plasticity can remain functional over long evolutionary time periods when the regulatory processes underlying plasticity are gradually broken down by mutations. To fix ideas, we consider a regulatory mechanism that switches on or off a pathway (like fat synthesis) in response to environmental conditions (e.g., host fat content).

#### Fitness considerations

We assume that the local environment of an individual is characterized by two factors: fat content *F* and nutrient content *N*, where nutrients represent sugars and other carbohydrates that can be used to synthesize fat. Nutrients are measured in units corresponding to the amount of fat that can be synthesized from them. We assume that fitness (viability and/or fecundity) is directly proportional to the amount of fat stored by the individual. When fat synthesis is switched off, this amount is equal to *F*, the amount of fat in the environment. When fat synthesis is switched on, the amount of fat stored is assumed to be *N* − *c*+(1−*k*)*F*. This expression reflects the following assumptions: *(i)* fat is synthesized from the available nutrients, but this comes at a fitness cost *c*; *(ii)* fat can still be absorbed from the environment, but at a reduced rate (1-*k*). It is adaptive to switch on fat synthesis if *N* − *c*+(1−*k*)*F* is larger than *F*, or equivalently if 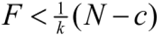.

The right-hand side of this inequality is a straight line, which is illustrated by the blue line in Fig. 4. The three boxes in Fig. 4 illustrate three types of environmental conditions.

**Fig. 4:**
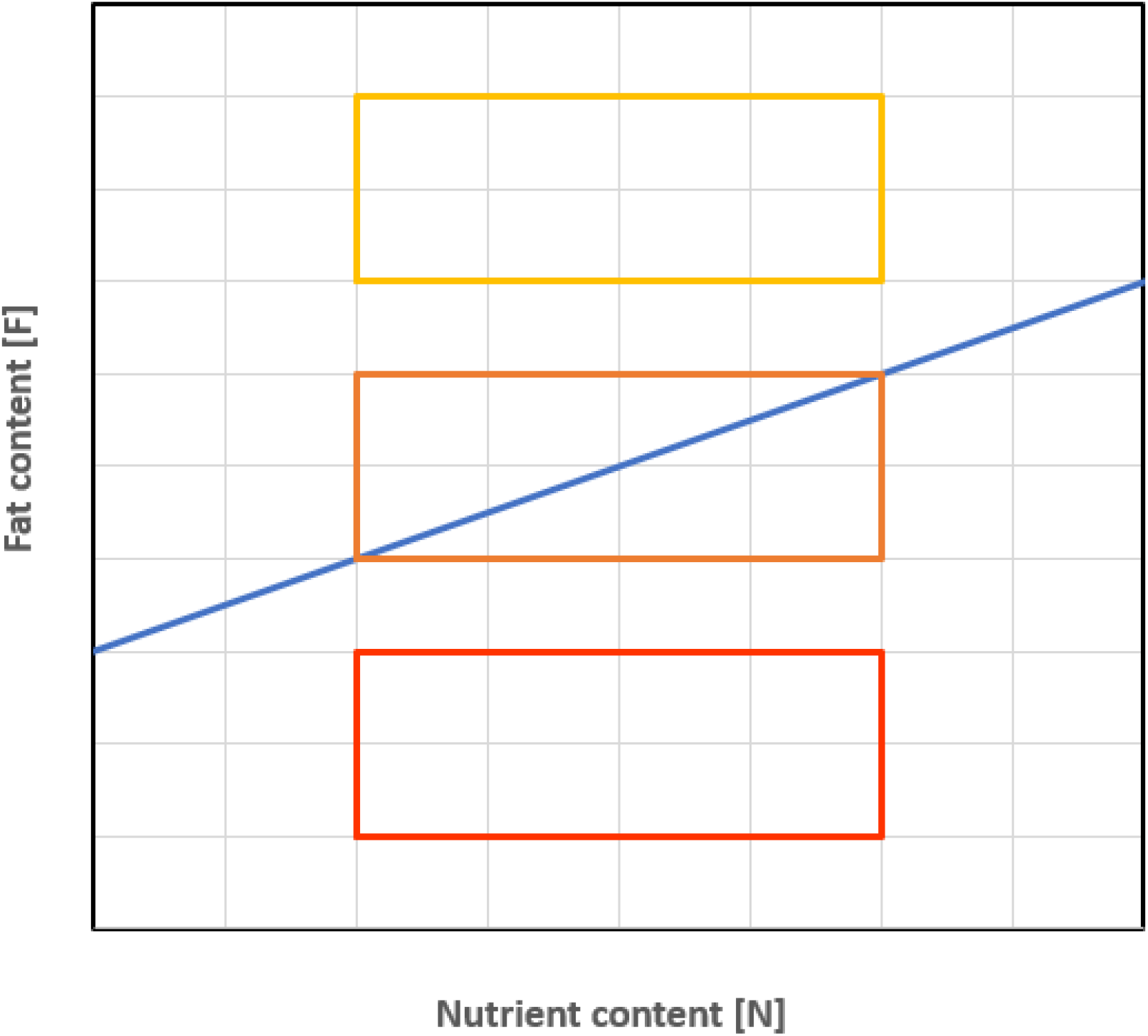
Environmental conditions encountered by the model organisms. For a given combination of environmental nutrient content *N* and environmental fat content *F*, it is adaptive to switch on fat synthesis if (*N*, *F*) is below the blue line (corresponding to 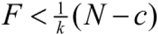) and to switch it off otherwise. The three boxes illustrate three types of environment: a low-fat environment (red) where fat synthesis should be switched on constitutively; a high-fat environment (yellow) where fat synthesis should be switched off constitutively; and an intermediate-fat environment (orange) where a plastic switch is selectively favoured.

- Red box: low-fat environments. Here, 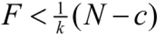 is always satisfied, implying that fat synthesis should be switched on constitutively.
- Yellow box: high-fat environments. Here, 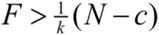, implying that fat synthesis should be switched off constitutively.
- Orange box: intermediate-fat environments. Here, fat synthesis should be plastic and switched on if for the given environment (*N*, *F*) the fat content is below the blue line and switched off otherwise.

The simulations reported here were all run for the parameters 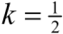 and 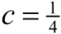.

#### Gene regulatory networks (GRN)

In our model, the switching device was implemented by an evolving gene regulatory network (as in^55^. The simulations shown in Fig. 3 of the main text are based on the simplest possible network that consists of two receptor nodes (sensing the fat and the nutrient content in the local environment, respectively) and an effector node that switches on fat synthesis if the combined weighted input of the two receptor nodes exceeds a threshold value *T* and switches it off otherwise. Hence, fat synthesis is switched on if *w*_*F*_*F*+ *w*_*N*_*N*>*T* (and off otherwise), where the weighing factors *w*_*F*_ and *w*_*N*_ and the threshold *T* are genetically determined evolvable parameters. We considered many alternative network structures (all with two receptor nodes and one effector node) and obtained very similar results (see below).

For the simple GRN described above, the switching device is 100% adaptive when the switch is on (i.e., *w*_*F*_*F* + *w*_*N*_*N*>*T*) if 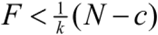 and off otherwise. A simple calculation yields that this is the case if: *w*_*N*_ > 0, *w*_*F*_ = −*kw*_*N*_ and *T* = *cw*_*N*_.

#### Evolution of the GRN

For simplicity, we consider an asexual haploid population with discrete, non-overlapping generations and fixed population size *N* = 10,000. Each individual has several gene loci, each locus encoding one parameter of the GRN. (In case of the simple network described above, there are three gene loci, which encode the parameters *w*_*F*_, *w*_*N*_ and *T*). At the start of its life, each individual is placed in a randomly chosen environment (*N*, *T*). Based on its (genetically encoded) GRN, the individual decides on whether to switch on or off fat synthesis. If synthesis is switched on, the individual’s fitness is given by *N*−*c*+(1−*k*)*F*; otherwise its fitness is given by *F*. Subsequently, the individuals produce offspring, where the number of offspring per individual is proportional to the individual’s fitness. Each offspring inherits the genetic parameters of its parents, subject to mutation. With probability μ (per locus) a mutation occurs. In such a case the parental value is changed by a certain amount; the mutational step size is drawn from a normal distribution with mean zero and standard deviation σ. In the reported simulations, we chose μ = 0.001 and σ = 0.1.

#### Preadaptation of the GRNs

Starting with randomly initialized population, we first let the population evolve in the intermediate-fat environment (orange box in Fig. 4) for 10,000 generations. In all replicate simulations, “perfectly adapted switch” (corresponding to *w*_*N*_ > 0, *w*_*F*_ = −*kw*_*N*_ and *T* = *cw*_*N*_) evolved, typically within 1,000 generations. These evolved networks were used to seed the populations in the subsequent “decay” simulations.

#### Evolutionary decay of the GRNs

For the decay experiments reported in Fig. 3 of the main text, we initiated a large number of monomorphic replicate populations with one of the perfectly adapted GRNs from the preadaptation phase. These populations were exposed for an extended period of time (1,000,000 generations) to a high-fat environment (yellow box in Fig. 4), where all GRNs switched off fat synthesis constitutively. However, in some scenarios, the environmental conditions changed back sporadically (with probability *q*) to the intermediate-fat environment, where it is adaptive to switch on fat metabolism in half of the environments. In Fig. 3, we report on the changing rates *q* = 0.0 (no changing back; red), *q* = 0.001 (changing back once every 1,000 generations; purple), and *q* = 0.01 (changing back once every 100 generations; pink). When such a change occurred, the population was exposed to the intermediate-fat environment for *t* generations (Fig. 3 is based on *t* = 3).

Throughout the simulation, the performance of the network was monitored every 100 generations as follows: 100 GRNs were chosen at random from the population, and each of these GRNs was exposed to 100 randomly chosen environmental conditions from the intermediate-fat environment. From this, we could determine the average percentage of “correct” decisions (where the network should be switched on if and only if 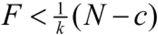. 1.0 means that the GRN is still making 100% adaptive decisions; 0.5 means that the GRN only makes 50% adaptive decision, as would be expected by a random GRN or a GRN that switches the pathway constitutively on or off. This measure for performance in the “old” intermediate-fat environment was determined for 100 replicate simulations per scenario and plotted in Fig. 3 (mean ± standard deviation).

#### Evolving robustness of the GRNs

The simulations in Fig. 3 are representative for all networks and parameters considered. Whenever *q* = 0.0, the performance of the regulatory switch eroded in evolutionary time, but typically at a much lower rate in case of the more complex GRNs. Whenever *q* = 0.01, the performance of the switch went back to levels above 90% and even above 95% for the more complex GRNs. Even for *q* = 0.001, a sustained performance level above 75% was obtained in all cases.

Intriguingly, in the last two scenarios the performance level first drops rapidly (from 1.0 to a much lower level, although this drop is less pronounced in the more complex GRNs) and subsequently recovers to reach high levels again. Apparently, the GRNs have evolved a higher level of robustness, a property that seems to be typical for evolving networks^8^. For the simple GRN studied in Fig. 3, this outcome can be explained as follows. The initial network was characterized by the genetic parameters *w*_*N*_ > 0, *w*_*F*_ = −*kw*_*N*_ and *T* = *cw*_*N*_ (see above), where *w*_*N*_ was typically a small positive number. In the course of evolutionary time, the relation between the three evolving parameters remained approximately the same, but *w*_*N*_ (and with it the other parameters) evolved to much larger values. This automatically resulted in an increasingly robust network, since mutations with a given step size distribution affect the performance of a network much less when the corresponding parameter is large in absolute value.

## Acknowledgements

We are grateful to Jerry Coyne, Frietson Galis, Georges Lognay, Philippe Vernon, Claude Remacle, René Rezsohazy, and David Denlinger for helpful suggestions on an earlier draft of this manuscript. We would like to thank Christophe Pels for maintaining hosts and parasitoids. This work was supported by the Fonds de la Recherche Scientifique - FNRS under grant n° 24905063 and 29109376. This is publication BRC 427 of the Biodiversity Research Centre.

## Author contributions

Conceptualization: BV, CMN; Formal analysis: BV, CMN; Modelling study: JMR, ST, TJBvE, FJW; Funding acquisition: BV, CMN, FJW; Investigation: BV, HTA, SR, MH; Resources: DR; Supervision: CMN, TH, FJW; Writing - original draft: BV, CMN, FJW; Writing - review & editing: all authors.

## Data availability

All data is available on https://visserlab.be

## References

1. Ellers, J., Kiers, T., Currie, C. R., Mcdonald, B. R. & Visser, B. Ecological interactions drive evolutionary loss of traits. Ecol. Lett. 15, 1071–1082 (2012).

2. Lahti, D. C. et al. Relaxed selection in the wild. Trends Ecol. Evol. 24, 487–496 (2009).

3. Collin, R. & Miglietta, M. P. Reversing opinions on Dollo’s law. Trends Ecol. Evol. 23, 602–609 (2008).

4. Esfeld, K. et al. Pseudogenization and resurrection of a speciation gene. Curr. Biol. 28, 3776–3786. (2019).

5. Zufall, R. A. & Rausher, M. D. Genetic changes associated with floral adaptation restrict future evolutionary potential. Nature 428, 847–850 (2004).

6. Tripp, E. A. & Manos, P. S. Is floral specialization an evolutionary dead-end? Pollination system transitions in Ruellia (Acanthaceae). Evolution (N. Y). 62, 1712–1737 (2008).

7. Lee, M. S. Y. & Shine, R. Reptilian viviparity and Dollo’s law. Evolution (N. Y). 52, 1441–1450 (1998).

8. Igic, B., Bohs, L. & Kohn, J. R. Ancient polymorphism reveals unidirectional breeding system shifts. Proc. Natl. Acad. Sci. U. S. A. 103, 1359–1363 (2006).

9. Domes, K., Norton, R. A., Maraun, M. & Scheu, S. Re-evolution of sexuality breaks Dollo’s Law. Proc. Natl. Acad. Sci. 104, 7139–7144 (2007).

10. Lynch, V. J. & Wagner, G. P. Did egg-laying boas break dollo’s law? Phylogenetic evidence for reversal to oviparity in sand boas (Eryx: Boidae). Evolution (N. Y). 64, 207–216 (2010).

11. Collin, R., Cipriani, R., Ambientales, D. D. E. & Simo, U. Dollo’s law and the re-evolution of shell coiling. Proc. R. Soc. B Biol. Sci. 270, 2551–2555 (2003).

12. Kohlsdorf, T. I. K. & Wagner, G. P. Evidence for the reversibility of digit loss: A phylogenetic study of limb evolution in *Bachia* (Gymnophthalmidae: Squamata). Evolution (N. Y). 60, 1896–1912 (2006).

13. Wiens, J. J. Re-evolution of lost mandibular teeth in frogs after more than 200 million years, and re-evaluating Dollo’s law. Evolution (N. Y). 65, 1283–1296 (2011).

14. Wiens, J. J. Re-evolution of lost mandibular teeth in frogs after more than 200 million years, and re-evaluating Dollo’s law. Evolution (N. Y). 65, 1283–1296 (2011).

15. Visser, B. & Ellers, J. Lack of lipogenesis in parasitoids: A review of physiological mechanisms and evolutionary implications. J. Insect Physiol. 54, 1315–1322 (2008).

16. Visser, B. et al. Loss of lipid synthesis as an evolutionary consequence of a parasitic lifestyle. Proc. Natl. Acad. Sci. 107, 8677–8682 (2010).

17. Turkish, A. R. & Sturley, S. L. The genetics of neutral lipid biosynthesis: An evolutionary perspective. Am. J. Physiol. Endocrinol. Metab. 297, E19–E27 (2009).

18. Jenke-kodama, H., Sandmann, A., Mu, R. & Dittmann, E. Evolutionary implications of bacterial polyketide synthases. Mol. Biol. Evol. 22, 2027–2039 (2003).

19. Maier, T., Leibundgut, M. & Ban, N. The crystal structure of a mammalian fatty acid synthase. Science (80-.). 321, 1315–1323 (2008).

20. Maier, T., Leibundgut, M., Boehringer, D. & Ban, N. Structure and function of eukaryotic fatty acid synthases. Q. Rev. Biophys. 43, 373–422 (2010).

21. Bukhari, H. S. T., Jakob, R. P. & Maier, T. Evolutionary origins of the multienzyme architecture of giant fungal fatty acid synthase. Struct. Des. 22, 1775–1785 (2014).

22. Peters, R. S. et al. Evolutionary history of the Hymenoptera. Curr. Biol. 27, 1013–1018 (2017).

23. Godfray, H. C. J. Parasitoids: Behavioural and evolutionary ecology. (Princeton University Press, 1994).

24. Prager, L., Bruckmann, A. & Ruther, J. *De novo* biosynthesis of fatty acids from α-D-glucose in parasitoid wasps of the *Nasonia* group. Insect Biochem. Mol. Biol. 115, 103256 (2019).

25. Visser, B. et al. Transcriptional changes associated with lack of lipid synthesis in parasitoids. Genome Biol. Evol. 4, 752–762 (2012).

26. Visser, B. et al. Variation in lipid synthesis, but genetic homogeneity, among Leptopilina parasitic wasp populations. Ecol. Evol. 8, 7355–7364 (2018).

27. Ament, S. A. et al. Mechanisms of stable lipid loss in a social insect. J. Exp. Biol. 214, 3808–21 (2011).

28. Visser, B. et al. Transcriptional changes associated with lack of lipid synthesis in parasitoids. Genome Biol. Evol. (2012).

29. Visser, B., Willett, D. S., Harvey, J. A. & Alborn, H. T. Concurrence in the ability for lipid synthesis between life stages in insects. R. Soc. Open Sci. 4, 160815 (2017).

30. Moiroux, J. et al. Local adaptations of life-history traits of a Drosophila parasitoid, Leptopilina boulardi: does climate drive evolution ? Ecol. Entomol. 35, 727–736 (2010).

31. Abu-Elheiga, L. et al. Mutant mice lacking acetyl-CoA carboxylase 1 are embryonically lethal. Proc. Natl. Acad. Sci. U. S. A. 102, 12011–12016 (2005).

32. Geer, B. W., Langevin, M. L. & McKechnie, S. W. Dietary ethanol and lipid synthesis in *Drosophila melanogaster*. Biochem. Genet. 23, 607–622 (1985).

33. Zinke, I., Schütz, C. S., Katzenberger, J. D., Bauer, M. & Pankratz, M. J. Nutrient control of gene expression in *Drosophila*: Microarray analysis of starvation and sugar-dependent response. EMBO J. 21, 6162–6173 (2002).

34. Wang, J. et al. Lipid dynamics, identification, and expression patterns of fatty acid synthase genes in an endoparasitoid, *Meteorus pulchricornis* (Hymenoptera: Braconidae). Int. J. Mol. Sci. 21, 1–14 (2020).

35. Wagner, A. Robustness and evolvability in living systems. (Princeton University Press, 2007).

36. Fleury, F., Gibert, P., Ris, N. & Allemand, R. Ecology and life history evolution of frugivorous *Drosophila*parasitoids. Adv. Parasitol. 70, 3–44 (2009).

37. Lue, C., Borowy, D., Buffington, M. L. & Leips, J. Geographic and seasonal variation in species diversity and community composition of frugivorous *Drosophila* (Diptera: Drosophilidae) and their *Leptopilina* (Hymenoptera: Figitidae) parasitoids. Environ. Entomol. 47, 1096–1106 (2018).

38. Hoffmann, A. R. Y. A. & Harshman, L. G. Desiccation and starvation resistance in *Drosophila*: patterns of variation at the species, population and intrapopulation levels. Heredity (Edinb). 83, 637–643 (1999).

39. Whiting, M. F., Bradler, S. & Maxwell, T. Loss and recovery of wings in stick insects. Nature 421, 264–267 (2003).

40. Domes, K., Norton, R. A., Maraun, M. & Scheu, S. Re-evolution of sexuality breaks Dollo’s Law. Proc. Natl. Acad. Sci. 104, 7139–7144 (2007).

41. Stone, G. & French, V. Evolution: Have wings come, gone and come again? Curr. Biol. 13, PR436–R438 (2003).

42. Goldberg, E. E. & Igic, B. On phylogenetic tests of irreversible evolution. Evolution (N. Y). 62, 2727–2741 (2008).

43. Christin, P.-A., Freckleton, R. P. & Osborne, C. P. Can phylogenetics identify C4 origins and reversals? Trends Ecol. Evol. 25, P403–P409 (2010).

44. Galis, F., Arntzen, J. W. & Lande, R. Dollo’s law and the irreversibility of digit loss in *Bachia*. Evolution (N. Y). 1–11 (2010).

45. Hall, B. K. Developmental mechanisms underlying the formation of atavisms. Biol. Rev. 59, 89–124 (1984).

46. Zhang, C.-X., Brisson, J. A. & Xu, H.-J. Molecular mechanisms of wing polymorphism in insects. Annu. Rev. Entomol. 64, 297–314 (2019).

47. Parker, D. J., Bast, J., Jalvingh, K. & Robinson-rechavi, M. Repeated evolution of asexuality involves convergent gene expression changes. Mol. Biol. Evol. 36, 350–364 (2018).

48. Tvedte, E. S., Jr, J. M. L. & Forbes, A. A. Sex loss in insects: causes of asexuality and consequences for genomes. Curr. Opin. Insect Sci. 31, 77–83 (2019).

49. Hanschen, E. R., Herron, M. D., Wiens, J. J., Nozaki, H. & Michod, R. E. Repeated evolution and reversibility of self-fertilization in the volvocine green algae. Evolution (N. Y). 72, 386–398 (2017).

50. R Development Core Team. R: A Language and Environment for Statistical Computing. (2016).

51. Kraaijeveld, K., Neleman, P., Marien, J., de Meijer, E. & Ellers, J. Genomic resources for *Goniozus legneri*, *Aleochara bilineata* and *Paykullia maculata*, representing three independent origins of the parasitoid lifestyle in insects. G3 Genes Genomes Genet. (2019).

52. Kraaijeveld, K. et al. Decay of sexual trait genes in an asexual parasitoid wasp. Genome Biol. Evol. 8, 3685–3695 (2016).

53. Marchler-Bauer, A. et al. CDD/SPARCLE: Functional classification of proteins via subfamily domain architectures. Nucleic Acids Res. 45, D200–D203 (2017).

54. Katoh, K. & Standley, D. M. MAFFT multiple sequence alignment software version 7: Improvements in performance and usability. Mol. Biol. Evol. 30, 772–780 (2013).

55. Gestel, J. Van & Weissing, F. J. Regulatory mechanisms link phenotypic plasticity to evolvability. Nat. Publ. Gr. 1–15 (2016).

